# Formation of templated inclusions in a forebrain α-synuclein mouse model is independent of LRRK2

**DOI:** 10.1101/2023.08.19.553965

**Authors:** Dylan J. Dues, Yue Ma, An Phu Tran Nguyen, Alina V. Offerman, Ian Beddows, Darren J. Moore

## Abstract

Leucine-rich repeat kinase 2 (LRRK2) and α-synuclein share enigmatic roles in the pathobiology of Parkinson’s disease (PD). *LRRK2* mutations are a common genetic cause of PD which, in addition to neurodegeneration, often present with abnormal deposits of α-synuclein in the form of Lewy-related pathology. As Lewy-related pathology is a prominent neuropathologic finding in sporadic PD, the relationship between LRRK2 and α-synuclein has garnered considerable interest. However, whether and how LRRK2 might influence the accumulation of Lewy-related pathology remains poorly understood. Through stereotactic injection of mouse α-synuclein pre-formed fibrils (PFF), we modeled the spread of Lewy-related pathology within forebrain regions where LRRK2 is most highly expressed. The impact of *LRRK2* genotype on the formation of α-synuclein inclusions was evaluated at 1-month post-injection. Neither deletion of *LRRK2* nor G2019S LRRK2 knockin appreciably altered the burden of α- synuclein pathology at this early timepoint. These observations fail to provide support for a robust pathophysiologic interaction between LRRK2 and α-synuclein in the forebrain *in vivo*. There was, however, a modest reduction in microglial activation induced by PFF delivery in the hippocampus of *LRRK2* knockout mice, suggesting that LRRK2 may contribute to α-synuclein-induced neuroinflammation. Collectively, our data indicate that the pathological accumulation of α-synuclein in the mouse forebrain is largely independent of LRRK2.

**Highlights:**

- Adult mice accumulate α-synuclein pathology in the hippocampus and cortex following stereotactic injection with α-synuclein PFFs, with negligible influence of *LRRK2* genotype.
- Hippocampal and cortical α-synuclein pathology elicits the concomitant accrual of phosphorylated tau, reactive astrogliosis, and microglial activation.
- Absence of endogenous *LRRK2* attenuates microglial activation in the dorsal hippocampus induced by PFFs, but not in the entorhinal cortex.
- Accumulation of α-synuclein inclusions and related neuropathologic changes were strongly associated across the hippocampal dorsal-ventral axis, regardless of *LRRK2* genotype.

## Introduction

Parkinson’s disease (PD) is a common neurodegenerative disorder defined by a complex constellation of clinical and pathological features (Langston, 2006). Patients with disease-causative mutations in the gene encoding leucine-rich repeat kinase 2 (LRRK2) account for one of the largest subgroups of familial PD (termed *LRRK2*-PD) (Biskup and West, 2008; Healy et al., 2008). Moreover, common genetic variation at the *LRRK2* locus has been implicated in PD susceptibility, suggesting that LRRK2 biology may be fundamental to both familial and sporadic disease (Nalls et al., 2019, 2014). LRRK2 pathogenicity is thought to arise from aberrantly enhanced kinase activity (West et al., 2007, 2005). There is also evidence to suggest that LRRK2 kinase activity may be elevated in cases of sporadic PD (Maio et al., 2018). Typical neuropathologic changes associated with sporadic PD are also detected in *LRRK2*-PD, including the accumulation of intraneuronal Lewy-related pathology. Thus, an understanding of whether LRRK2 impacts the accumulation of Lewy-related pathology, a major neuropathological hallmark of PD, is of critical interest.

The neuropathology of *LRRK2*-PD is diverse (Paisán-Ruı z et al., 2004; Zimprich et al., 2004). Degeneration of the dopaminergic nigrostriatal system remains a unifying feature and necessary element for slowness of movement. However, pathologic heterogeneity has been appreciated in almost all post-mortem studies to date (Wider et al., 2010). Lewy-related pathology has been identified as the prominent histopathologic feature in most cases, though other proteinopathies have been detected in combination with or in lieu of Lewy-related pathology (Schneider and Alcalay, 2017). Interestingly, some *LRRK2*-PD cases exhibit “pure” neurodegeneration in the absence of protein aggregation (Hasegawa et al., 2009; Takanashi et al., 2018). The lack of Lewy-related pathology in a substantial number of *LRRK2*-PD cases has spawned interest in alternative pathological substrates. This exploration of other proteinopathies has led to recent identification of AD-like neurofibrillary tangles, amyloid-beta plaques, and TDP- 43 inclusions in cases of *LRRK2*-PD. Towards this, a recent case series found that AD-like tau pathology was ubiquitous within a small cohort of *LRRK2*-PD cases (Henderson et al., 2019c). Beyond protein aggregation, several other neuropathological changes are detected in both sporadic PD and *LRRK2*-PD including astroglial reactivity and microglial activation.

While the neuropathology of *LRRK2*-PD is pleiotropic, it is unclear how various proteinopathies might arise in the context of pathogenic LRRK2 activity. In any event, the frequency with which *LRRK2*-PD cases exhibit Lewy-related pathology has led to the query of whether LRRK2 might modulate the pathologic accumulation of α- synuclein, the major proteinaceous component of Lewy bodies. A range of experimental studies have explored this query in animal models, with mixed results (Dues and Moore, 2020). Given the development of LRRK2-targeted therapeutics, the impact of such approaches on the progression of Lewy-related pathology remains uncertain. This is concerning, as the accumulation of Lewy-related pathology within forebrain structures has been associated with cognitive dysfunction and dementia in PD (Aarsland et al., 2005). As nearly 80% of PD patients will develop dementia, the contributions of LRRK2 towards the expansion of Lewy-related pathology within the forebrain require clarification (Hely et al., 2008). Indeed, LRRK2-targeted therapeutics may provide a critical approach to disease-modification if the accumulation of Lewy-related pathology in the forebrain can be halted.

Here, we aim to determine whether LRRK2 modifies the accumulation of Lewy-like α-synuclein inclusions within the hippocampus and cortex of an α-synuclein pre-formed fibril (PFF) mouse model. We opted to use this approach provided that α-synuclein PFFs are a well-established tool for generating inclusions *in vivo* (Duffy et al., 2018). Moreover, the application of α-synuclein PFFs to the forebrain has been demonstrated to elicit hippocampal neurodegeneration and cognitive dysfunction in mice, of relevance to Lewy-related cognitive dysfunction (Dues et al., 2023). We unilaterally injected α- synuclein PFFs into the forebrain and assessed the burden of α-synuclein pathology at 1-month post-injection (1 MPI) within two separate cohorts (wild-type vs G2019S knockin; wild-type vs LRRK2 knockout). Our results clearly demonstrate that the presence of endogenous LRRK2 is not an obligate requirement for the formation of α- synuclein inclusions within the mouse forebrain, like previous observations in transgenic α-synuclein models (Daher et al., 2012). Moreover, *LRRK2* genotype did not significantly alter the burden of α-synuclein pathology (as measured by pS129-α- synuclein immunostaining) across hippocampal and cortical regions. Several related neuropathological changes, including markers of microglial and astroglial reactivity, were found to be elevated in inclusion-bearing regions. Intriguingly, the absence of endogenous LRRK2 revealed a modest reduction in microglial activation within the dorsal hippocampus. Collectively, our results find little evidence of a substantive contribution of LRRK2 towards templated α-synuclein pathology in the mouse forebrain.

## Results

### Stereotactic injection of α-synuclein PFFs into the mouse forebrain

Our experimental design sought to assess the impact of endogenous LRRK2 in comparison to either G2019S LRRK2 or *LRRK2* deletion on the accumulation of α- synuclein pathology in adult mice (**Figure 1A, B**). The G2019S mutation is one of the most frequently detected mutations causative for *LRRK2*-PD (Guedes et al., 2010). We opted to utilize an intra-hippocampal/cortical injection paradigm with α-synuclein PFFs to trigger the robust formation of templated α-synuclein inclusions in the mouse forebrain (Dues et al., 2023). LRRK2 expression in these regions is well-established (Daher et al., 2012; West et al., 2014). To confirm the cellular localization of endogenous *LRRK2* expression in the mouse forebrain, we utilized *in situ* hybridization combined with immunofluorescence. RNAScope probes directed against mouse *LRRK2* mRNA were used to identify *LRRK2* transcript in hippocampal neurons (**Figure 2**). Consistent with previous studies, we found that expression was abundant in the hippocampus and cortex (Daher et al., 2012) Expression was generally localized to NeuN-positive neurons rather than in other non-neuronal cell-types, as anticipated (**Figure 2**).

**Figure 1.**
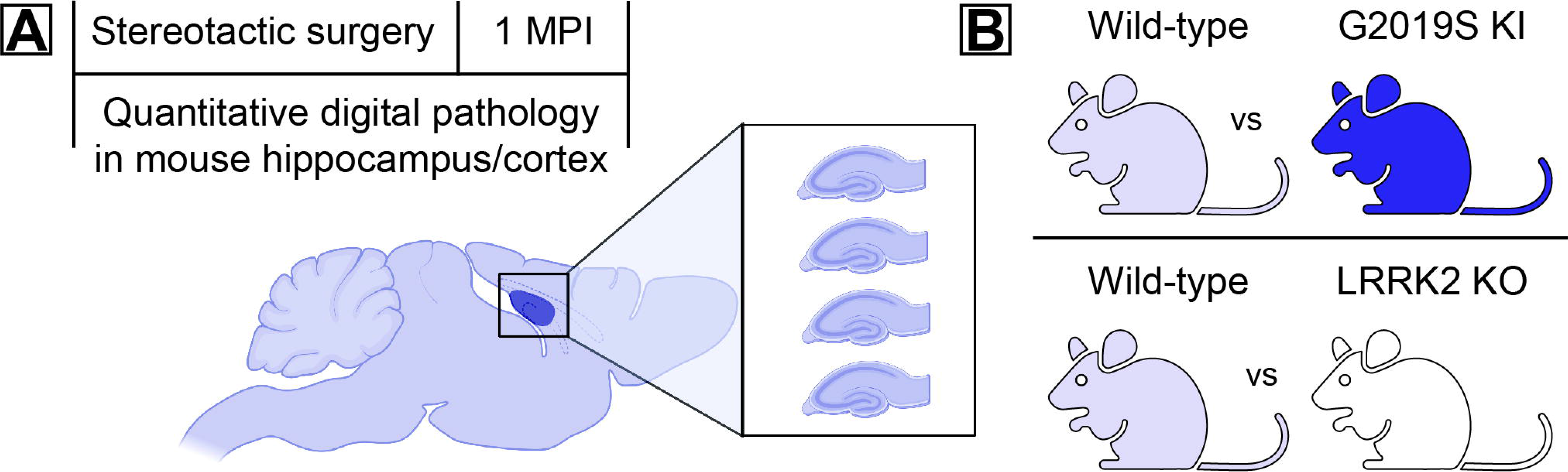
Experimental design. **A)** Adult mice (C57BL/6J) were subjected to unilateral stereotactic injection into the forebrain with α-synuclein PFFs. Cohorts were assessed at 1 MPI and serial horizontal sections of hippocampus and cortex were assessed using a digital pathology approach. **B)** Cohorts were composed of either wild-type and G2019S LRRK2 knockin (KI) mice or wild-type and LRRK2 knockout (KO) mice.

**Figure 2.**
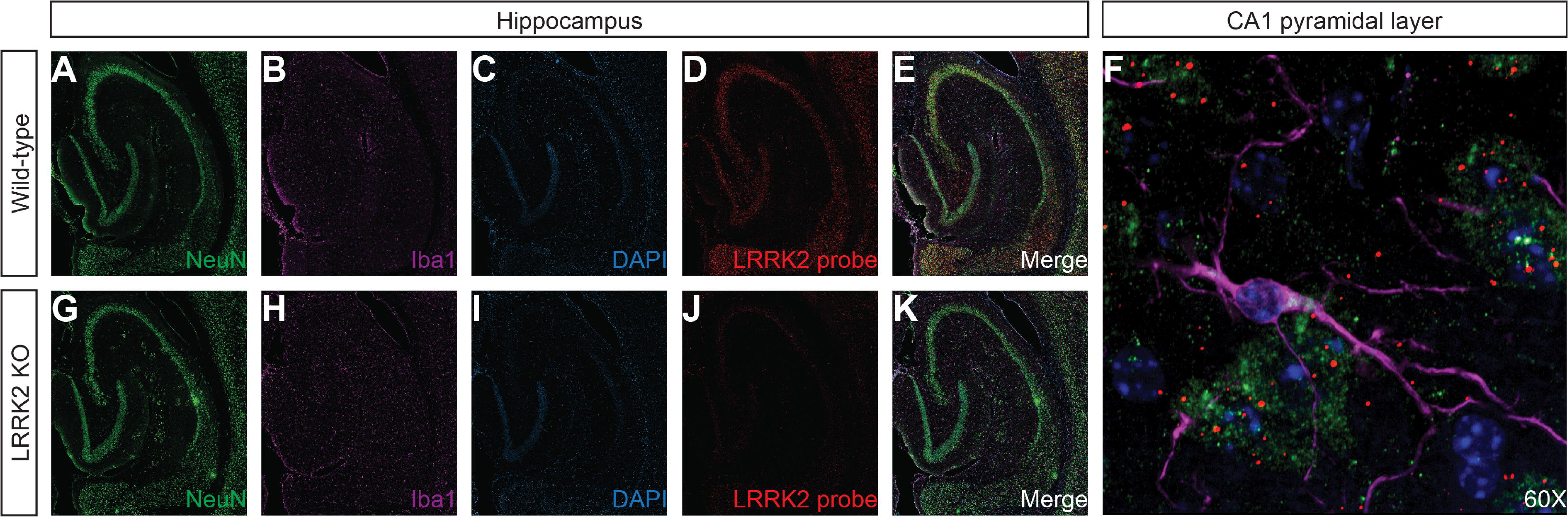
Localization of LRRK2 expression in the mouse hippocampus. **A-F)** Representative hippocampal section from a wild-type mouse displaying various markers including **(A)** NeuN, **(B)** Iba1, **(C)** DAPI, **(D)** LRRK2 RNAScope probe, and **(E)** merged. A 60x zoomed image **(F)** of the previous merged images exhibiting LRRK2 expression in NeuN-positive neurons but not in Iba1-positive microglia. **G-K)** Representative hippocampal section from a LRRK2 knockout mouse displaying various markers including (**G**) NeuN, **(H)** Iba1, **(I)** DAPI, **(J)** absence of LRRK2 RNAScope probe signal, and **(K)** merged.

### The burden of **α**-synuclein pathology in the hippocampus and cortex is not substantially altered by LRRK2

Following the unilateral injection of α-synuclein PFFs into the mouse forebrain, we evaluated the distribution and burden of α-synuclein pathology. Inclusions were detected by pS129-α-synuclein immunostaining. Pathologic α-synuclein was distributed throughout the hippocampal dorsal-ventral axis, with the bulk of pathology accumulating towards the dorsal portion of the hippocampus (**Figure 3A, 4A**). We also observed the spread of α-synuclein pathology to the hemisphere contralateral to the injection site (**Figure 3A, 4A**). Of note, the burden of hippocampal pathology was quite substantial in either hemisphere due to extensive inter-hemispheric connectivity. In contrast, the formation of pathology in the cortex was effectively limited to the hemisphere ipsilateral to the injection site. Specifically, the ipsilateral cortex exhibited robust α-synuclein pathology in layers II/III of the lateral entorhinal cortex (**Figure 3F, 4F**). Mice expressing G2019S LRRK2 did not exhibit a regional distribution or burden of α-synuclein pathology that was substantially different from wild-type mice (**Figure 3B-E**). This finding was consistent in both the hippocampus and cortex. Likewise, deletion of endogenous *LRRK2* also failed to alter the burden of α-synuclein pathology (**Figure 4B- E**). In *LRRK2* knockout mice, α-synuclein inclusions readily formed with a tendril-like morphology indistinguishable from that of wild-type mice (**Figure 4F**). Taken together, these findings support that LRRK2 and α-synuclein are unlikely to share a direct pathophysiologic interaction in neurons of the forebrain. Further, it appears unlikely that the *in vivo* accumulation of α-synuclein pathology is substantially modified by LRRK2.

**Figure 3.**
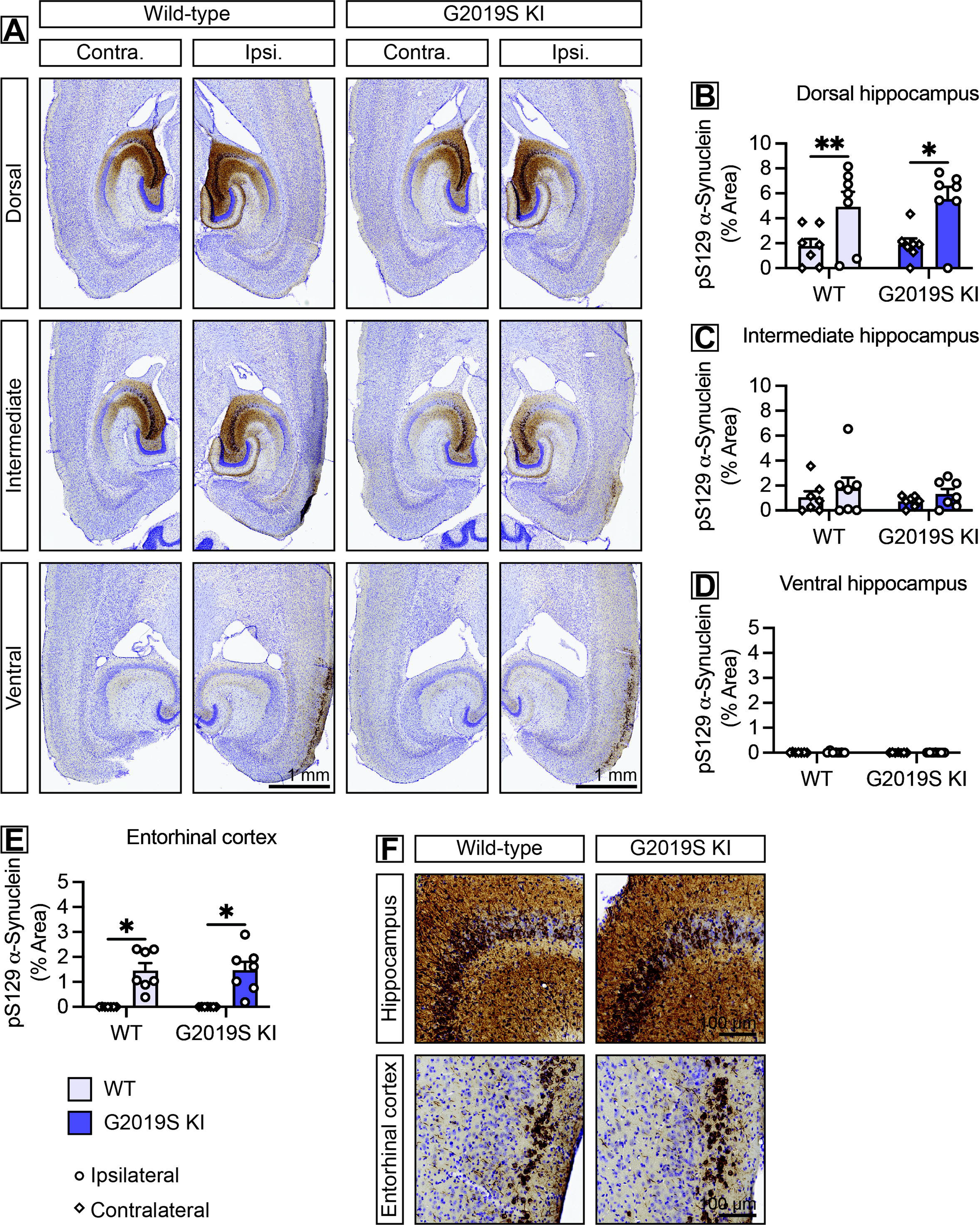
G2019S LRRK2 expression does not augment hippocampal or cortical α- synuclein pathology. **A)** Representative histological sections from wild-type and *G2019S LRRK2* knockin (KI) mice spanning the hippocampal axis and immunostained for pS129-α-synuclein. Ipsilateral (Ipsi.) refers to the injected hemisphere while contralateral (Contra.) refers to the non-injected hemisphere. Scale bars: 1 mm. **B-E)** Quantification of pS129-α- synuclein immunostaining by % area across the **(B)** dorsal, **(C)** intermediate, and **(D)** ventral portions of the hippocampus and in the **(E)** entorhinal cortex. Data are expressed as bars depicting the mean ± SEM from either the ipsilateral or the contralateral hemisphere with each data point representing an animal (*n* = 7 animals/genotype). \**P*<0.05 or \*\**P*<0.01 by paired *t*-test or Wilcoxon matched-pairs signed rank test within genotype, as indicated. Non-significant by unpaired *t*-test or Mann Whitney test between genotypes. **F)** Representative histological sections at higher magnification depicting hippocampal and cortical pS129-α-synuclein immunostaining. Scale bars: 100 μm.

**Figure 4.**
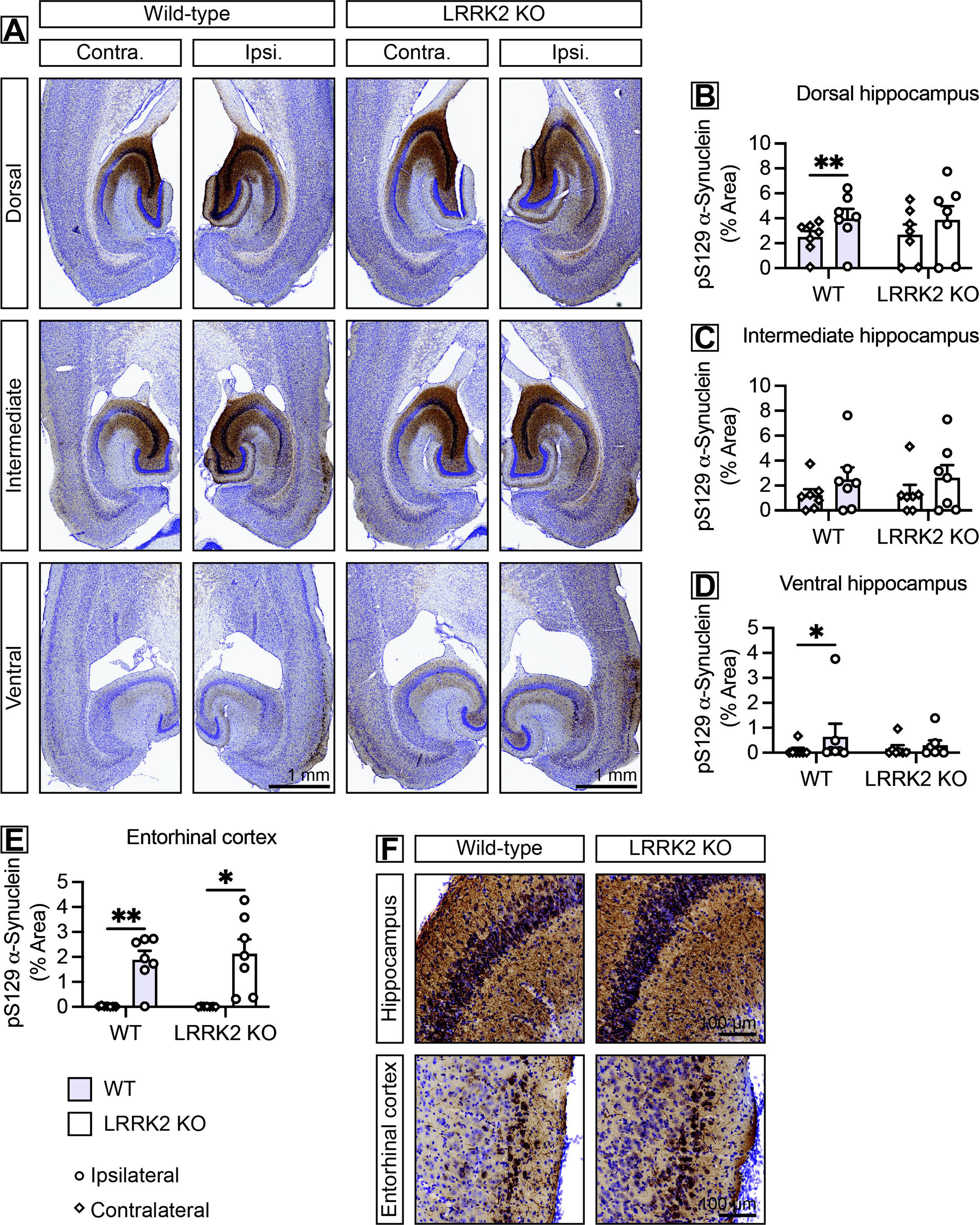
Absence of endogenous LRRK2 does not reduce hippocampal or cortical α-synuclein pathology. **A)** Representative histological sections from wild-type and *LRRK2* knockout (KO) mice spanning the hippocampal axis and immunostained for pS129-α-synuclein. Ipsilateral (Ipsi.) refers to the injected hemisphere while contralateral (Contra.) refers to the non-injected hemisphere. Scale bars: 1 mm. **B-E)** Quantification of pS129-α-synuclein immunostaining by % area across the **(B)** dorsal, **(C)** intermediate, and **(D)** ventral portions of the hippocampus and in the **(E)** entorhinal cortex. Data are expressed as bars depicting the mean ± SEM from either the ipsilateral or the contralateral hemisphere with each data point representing an animal (*n* = 7 animals/genotype). \**P*<0.05 or \*\**P*<0.01 by paired *t*-test or Wilcoxon matched-pairs signed rank test within genotype, as indicated. Non-significant by unpaired *t*-test or Mann Whitney test between genotypes. **F)** Representative histological sections at higher magnification depicting hippocampal and cortical pS129-α-synuclein immunostaining. Scale bars: 100 μm.

### Accumulation of phosphorylated tau granules occurs following injection with **α**-synuclein PFFs

An intriguing feature of the α-synuclein PFF model is that PFF-injection has previously been found to lead to the aberrant mislocalization and accumulation of tau *in vivo* (Guo et al., 2013). A distinct strain of α-synuclein fibrils has been shown to induce AD-like neurofibrillary tangle pathology in mice (Guo et al., 2013). Further, several groups have found that α-synuclein influences the spread of tau pathology (Bassil et al., 2020; Williams et al., 2020). While it is unclear if this cascade results from a direct cross-seeding event or rather as a conditional response to neuronal proteotoxic stress or axonal injury, the concomitant accumulation of tau in our model is likely relevant to *LRRK2*-PD pathobiology. *LRRK2*-PD is known to exhibit pathological accumulation of AD-like tau pathology, and several LRRK2 models have exposed a potential interaction between LRRK2 and tau (Bailey et al., 2013; Cornblath et al., 2021; Nguyen et al., 2018). Thus, we sought to evaluate whether the subsequent accumulation of phosphorylated tau granules in our model was influenced by LRRK2.

We identified extensive accumulation of phosphorylated tau (AT8-positive) granules in both hemispheres of the hippocampus, as well as in the entorhinal cortex, in parallel with the accumulation of α-synuclein pathology (**Figure 5E, J**). Intriguingly, these tau granules appeared to localize within neuronal processes, though smaller puncta appeared to cluster within cell bodies. Provided that the entorhinal cortex is proposed to be especially vulnerable to neurofibrillary tangle formation, it is intriguing that tau accumulation in this region appear to localize more prominently within cell bodies. This divergence in cellular localization may be related to specific neuronal vulnerabilities to either α-synuclein or tau. In either case, we found that endogenous LRRK2 is not required for the formation of these granules, and that the burden was not appreciably altered by *LRRK2* genotype (**Figure 5A-D, F-I**).

**Figure 5.**
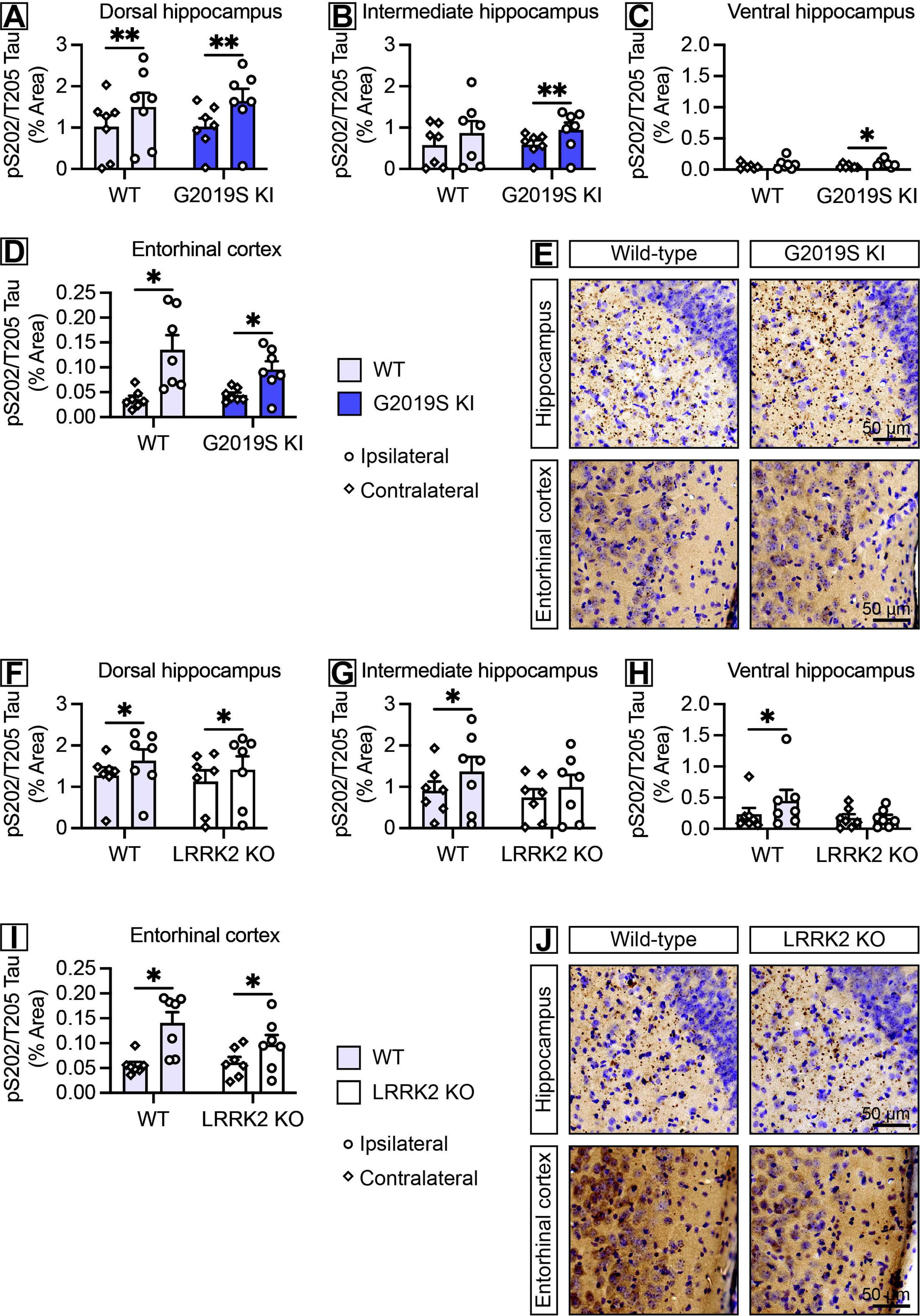
Accumulation of AT8-positive tau is not altered by *LRRK2* genotype. **A-D)** Quantification of pS202/T205-tau immunostaining by % area across the **(A)** dorsal, **(B)** intermediate, and **(C)** ventral portions of the hippocampus and in the **(D)** entorhinal cortex of wild-type and G2019S LRRK2 knockin mice. Data are expressed as bars depicting the mean ± SEM from either the ipsilateral or the contralateral hemisphere with each data point representing an animal (n = 7 animals/genotype). *P<0.05 or **P<0.01 by paired t-test or Wilcoxon matched-pairs signed rank test within genotype, as indicated. Non-significant by unpaired t-test or Mann Whitney test between genotypes. **E)** Representative histological sections depicting hippocampal and cortical pS202/T205-tau immunostaining. Scale bars: 50 μm. **F-I)** Quantification of pS202/T205-tau immunostaining by % area across the **(F)** dorsal, **(G)** intermediate, and **(H)** ventral portions of the hippocampus and in the **(I)** entorhinal cortex of wild-type and LRRK2 knockout mice. Data are expressed as bars depicting the mean ± SEM from either the ipsilateral or the contralateral hemisphere with each data point representing an animal (n = 7 animals/genotype). *P<0.05 by paired t-test or Wilcoxon matched-pairs signed rank test within genotype, as indicated. Non-significant by unpaired t-test or Mann Whitney test between genotypes. **J)** Representative histological sections depicting hippocampal and cortical pS202/T205-tau immunostaining. Scale bars: 50 μm.

### Astroglial reactivity following the accumulation of **α**-synuclein pathology is not altered by *LRRK2* genotype

Astrogliosis is a hallmark feature of PD and is associated with neurodegenerative changes in a variety of experimental models (Booth et al., 2017). We sought to determine whether astroglial reactivity, in response to α-synuclein pathology, would be modified by *LRRK2* genotype. We noted marked elevation in GFAP-positive immunostaining in portions of the hippocampus and cortex affected by α-synuclein pathology (**Figure 6E, J**). Comparable with regional burdens of α-synuclein pathology, we found elevated levels of astrogliosis in both hemispheres of the hippocampus, while the entorhinal cortex exhibited relatively higher levels of astrogliosis on the ipsilateral hemisphere. As with α-synuclein pathology, we found no impact of *LRRK2* genotype on astroglial activity as measured by GFAP levels (**Figure 6A-D, F-I**).

**Figure 6.**
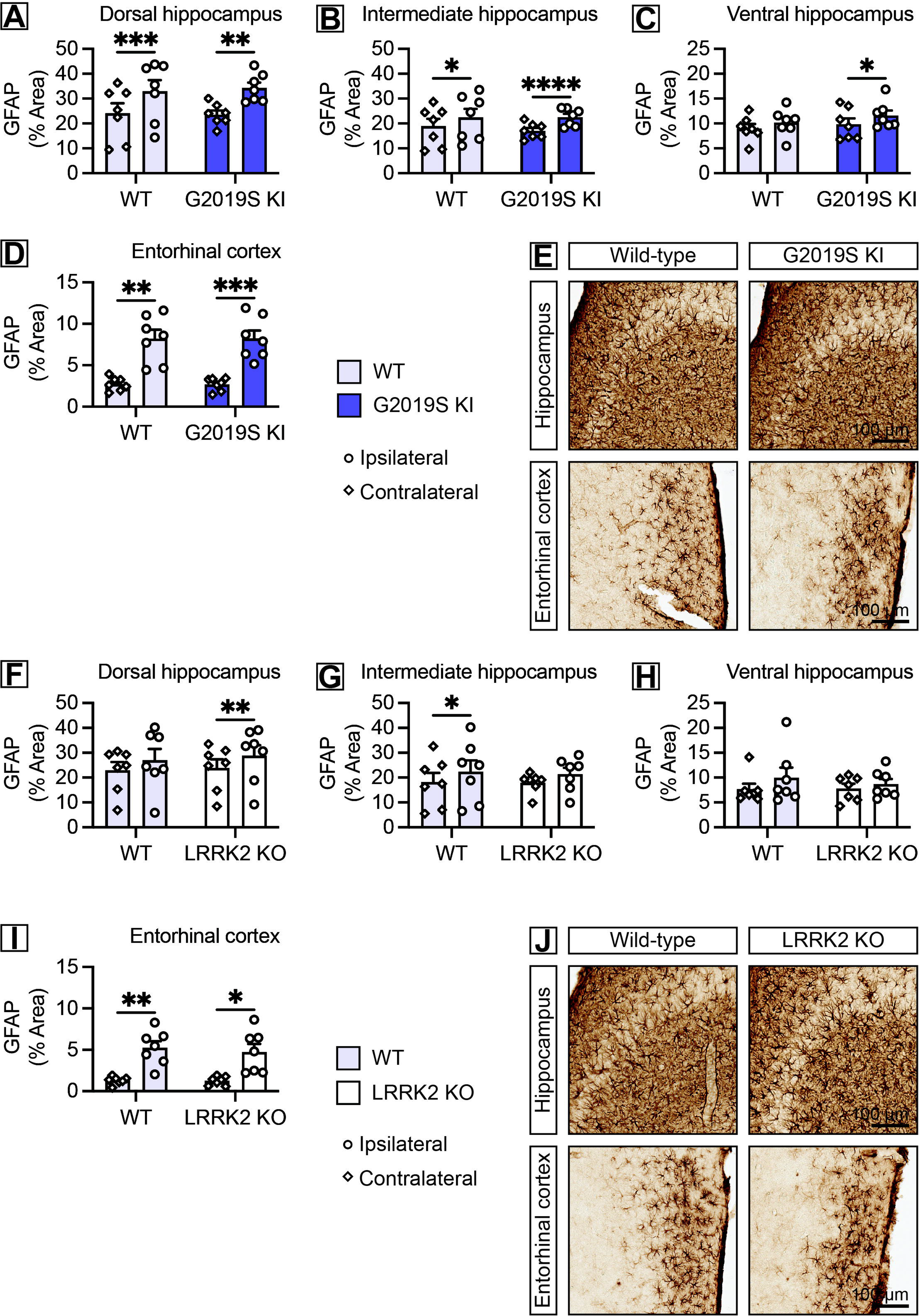
Astroglial reactivity is not impacted by *LRRK2* genotype. **A-D)** Quantification of GFAP immunostaining by % area across the **(A)** dorsal, **(B)** intermediate, and **(C)** ventral portions of the hippocampus and in the **(D)** entorhinal cortex of wild-type and *G2019S LRRK2* knockin mice. Data are expressed as bars depicting the mean ± SEM from either the ipsilateral or the contralateral hemisphere with each data point representing an animal (*n* = 7 animals/genotype). \**P*<0.05, \*\**P*<0.01, \*\*\**P*<0.001, or \*\*\*\**P*<0.0001 by paired *t*-test or Wilcoxon matched-pairs signed rank test within genotype, as indicated. Non-significant by unpaired *t*-test or Mann Whitney test between genotypes. **E)** Representative histological sections depicting hippocampal and cortical GFAP immunostaining. Scale bars: 100 μm. **F-I)** Quantification of GFAP immunostaining by % area across the **(F)** dorsal, **(G)** intermediate, and **(H)** ventral portions of the hippocampus and in the **(I)** entorhinal cortex of wild-type and *LRRK2* knockout mice. Data are expressed as bars depicting the mean ± SEM from either the ipsilateral or the contralateral hemisphere with each data point representing an animal (*n* = 7 animals/genotype). \**P*<0.05 or \*\**P*<0.01 by paired *t*-test or Wilcoxon matched-pairs signed rank test within genotype, as indicated. Non-significant by unpaired *t*-test or Mann Whitney test between genotypes. **J)** Representative histological sections depicting hippocampal and cortical GFAP immunostaining. Scale bars: 100 μm.

### Microglial activation is modestly dampened in the hippocampus of *LRRK2* knockout mice

Previous studies have illuminated a role for microglia in the pathogenesis of PD (Imamura et al., 2003; Kannarkat et al., 2013). LRRK2 is speculated to influence inflammatory changes, including the microglial response to various stimuli (Gillardon et al., 2012; Kim et al., 2012; Marker et al., 2012; Moehle et al., 2012). However, there is limited evidence of LRRK2 protein within microglia of the mouse brain, despite evidence of LRRK2 activation in mouse primary microglial cultures and in human microglia (Kozina et al., 2018; Langston et al., 2022). As expected, we observed amoeboid-like changes in microglial morphology around pathology-affected regions (detected by Iba1 immunostaining) in mice injected with α-synuclein PFFs (**Figure 7E, J**). Thus, we examined whether *LRRK2* genotype might alter the response of microglia in our model. Comparable with the distribution of α-synuclein pathology, we observed altered Iba1-positive immunostaining in the dorsal hippocampus and cortex. As proxy for activation, we observed enlarged microglial cell bodies with retracted processes, consistent with an amoeboid-like morphology. We did not detect a difference in Iba1 immunostaining between wild-type and G2019S LRRK2 knockin mice, suggesting that pathogenic LRRK2 expression does not augment the microglial response to α-synuclein pathology (**Figure 7A-D**). In contrast, we detected a modest reduction in Iba1-positive immunostaining in the dorsal portion of the hippocampus in *LRRK2* knockout mice (**Figure 7F)**. Other hippocampal regions, as well as the entorhinal cortex, did not exhibit any differences in total Iba1 levels **(Figure 7G-I)**.

**Figure 7.**
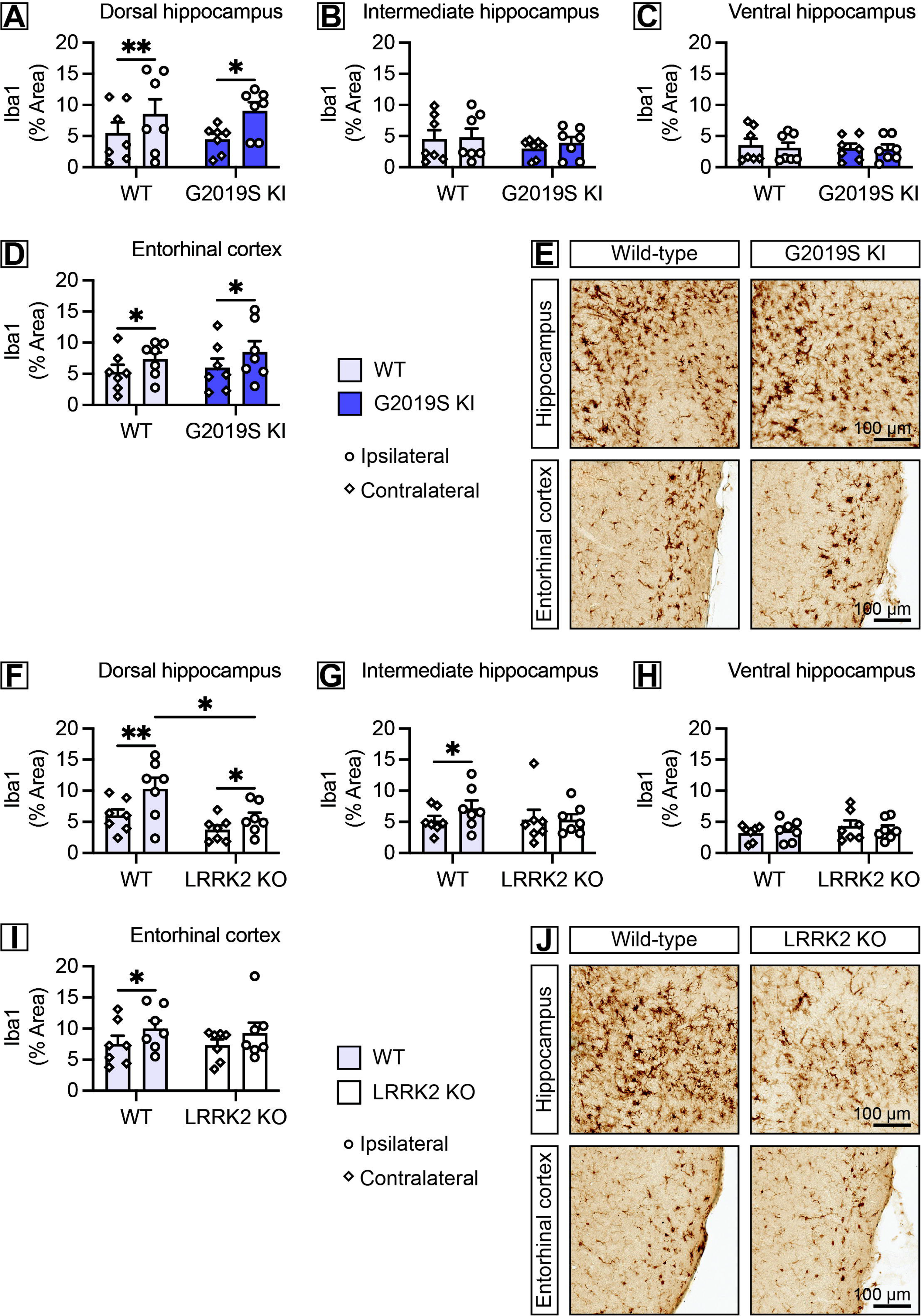
*LRRK2* knockout mice exhibit a modest reduction in microglial activation in the hippocampus. **A-D)** Quantification of Iba1 immunostaining by % area across the **(A)** dorsal, **(B)** intermediate, and **(C)** ventral portions of the hippocampus and in the **(D)** entorhinal cortex of wild-type and *G2019S LRRK2* knockin mice. Data are expressed as bars depicting the mean ± SEM from either the ipsilateral or the contralateral hemisphere with each data point representing an animal (*n* = 7 animals/genotype). \**P*<0.05 or \*\**P*<0.01 by paired *t*-test or Wilcoxon matched-pairs signed rank test within genotype, as indicated. Non-significant by unpaired *t*-test or Mann Whitney test between genotypes. **E)** Representative histological sections depicting hippocampal and cortical Iba1 immunostaining. Scale bars: 100 μm. **F-I)** Quantification of Iba1 immunostaining by % area across the **(F)** dorsal, **(G)** intermediate, and **(H)** ventral portions of the hippocampus and in the **(I)** entorhinal cortex of wild-type and *LRRK2* knockout mice. Data are expressed as bars depicting the mean ± SEM from either the ipsilateral or the contralateral hemisphere with each data point representing an animal (*n* = 7 animals/genotype). \**P*<0.05 or \*\**P*<0.01 by paired *t*-test or Wilcoxon matched-pairs signed rank test within genotype, as indicated. \**P*<0.05 by unpaired *t*-test between genotypes, as indicated. **J)** Representative histological sections depicting hippocampal and cortical Iba1 immunostaining. Scale bars: 100 μm.

Importantly, Iba1 is known to be expressed by both resident microglia as well as peripherally recruited macrophages. To clarify whether α-synuclein pathology (or α-synuclein PFF) led to the recruitment of peripheral immune cells, we conducted additional immunostaining for CD45 (**Figure S1A**). CD45 expression is highly elevated in phagocytic cells recruited from the periphery. We observed the recruitment of CD45-positive peripheral immune cells into the dorsal hippocampus, but not in other regions. Quantitation revealed a trend towards a reduction of CD45-positive cell counts in *LRRK2* knockout mice, but this was not significant, and counts in G2019S LRRK2 knockin mice did not differ substantially from wild-type mice (**Figure S1B, C**). Notably, other groups have also identified peripheral immune cell recruitment following induction of α-synuclein pathology (Harms et al., 2018; Xu et al., 2022). It is plausible that LRRK2 activation may influence this process, and mutant LRRK2 expression has been demonstrated to impact myeloid cell chemotaxis (Moehle et al., 2015).

### Microglial morphologic changes are detected in the entorhinal cortex but are not substantially altered by LRRK2

To explore whether LRRK2 might influence the local microglial response to intraneuronal α-synuclein pathology, we sought to assess microglial morphology in the entorhinal cortex. The ipsilateral entorhinal cortex exhibits robust α-synuclein pathology in this model and is located distal to the injection site, limiting direct exposure to extracellular fibrils. Notably, the hippocampus is the only region in which resident microglia would predictably be exposed to both extracellular PFF as well as intraneuronal pathology. In contrast, microglia in the entorhinal cortex would seemingly be provoked in response to intraneuronal α-synuclein without exposure to extracellular PFF. Thus, the region-specific basis for this dampening in Iba1-positive immunostaining in the dorsal hippocampus might be explained by LRRK2 influencing the response to extracellular fibrillar α-synuclein rather than as a response to intraneuronal inclusions. Microglia have previously been demonstrated to react to fibrillar α-synuclein *in vitro*, so it is plausible that the absence of LRRK2 activity has a deleterious effect on this aspect of microglial biology (Kim et al., 2020; Moehle et al., 2012). Alternatively, the increased density of resident microglia within the hippocampus, or greater abundance of pathological α-synuclein at this site may explain the dichotomy of microglial activation in the hippocampus and cortex.

We observed clear enlargement of the cell body in some Iba1-positive cells in the ipsilateral entorhinal cortex and utilized a higher threshold gradient of optical density to identify strong immunostained cells (**Figure 8C**). Using this more restrictive threshold, we detected significantly elevated Iba1 immunostaining in the ipsilateral entorhinal cortex of all experimental groups except for *LRRK2* knockout mice (**Figure 8A, B**). To conduct a more detailed analysis of altered microglial morphology in the entorhinal cortex, we employed a microglial activation module from the HALO digital pathology platform to identify and quantify the cell body size of individual microglia (**Figure 8H**). We observed a clear shift in the distribution of microglia cell body size in the ipsilateral entorhinal cortex, consistent with microglial activation in this region (**Figure 8F, G**). The mean Iba1-positive cell body area was determined to be significantly increased on the ipsilateral entorhinal cortex relative to the contralateral entorhinal cortex, indicative of pathology-associated activation of microglia. Intriguingly, a modest but significant increase in mean microglial cell body size was detected in the contralateral entorhinal cortex of *LRRK2* knockout mice (**Figure 8D, E**). While further analysis is necessary, this may be related to an intrinsic alteration of microglial function in *LRRK2* knockout mice.

**Figure 8.**
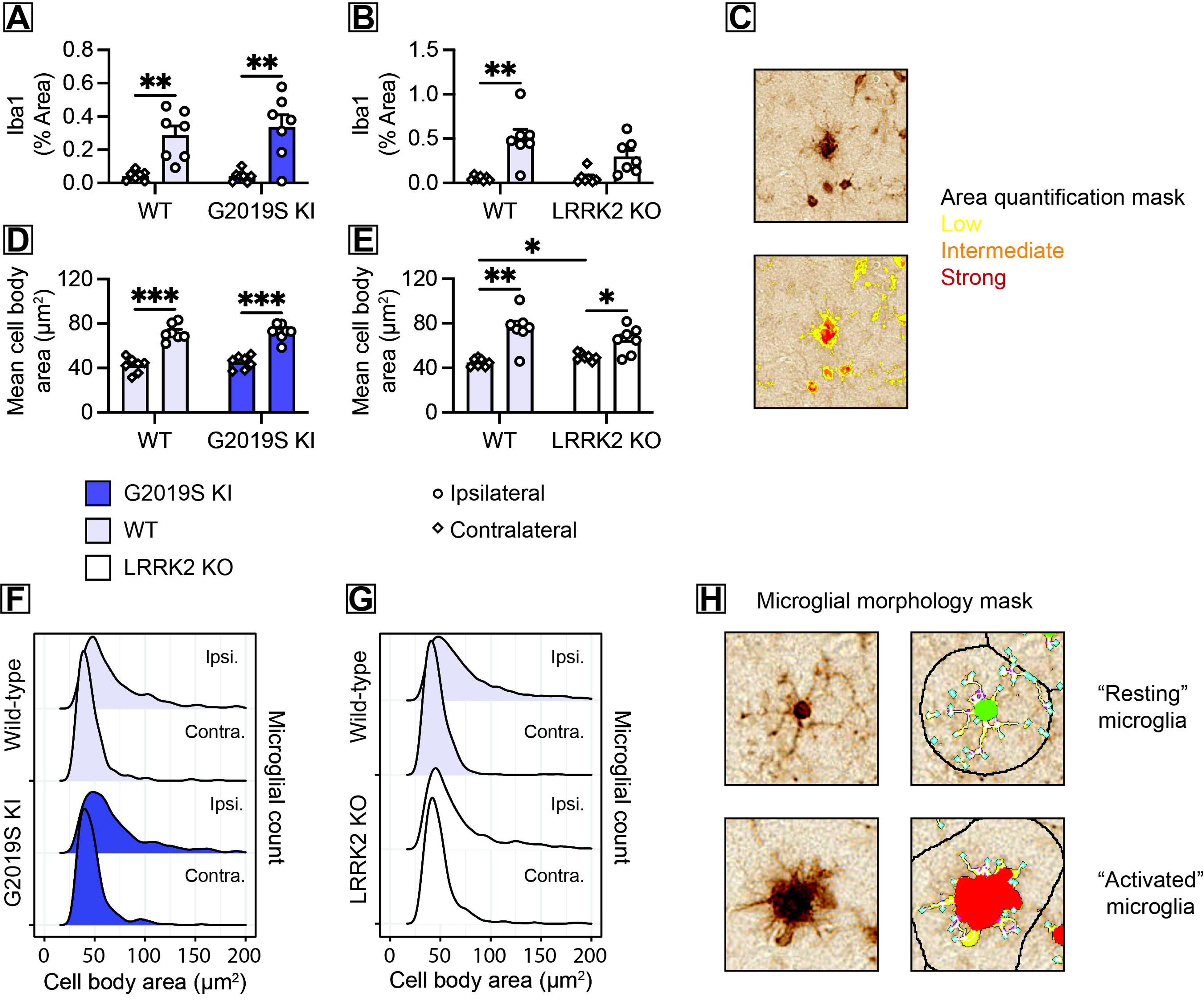
The formation of α-synuclein inclusions in the entorhinal cortex leads to substantial alterations in microglial morphology independent of *LRRK2* genotype. **A-B)** Quantification of Iba1 immunostaining by % area for “strong” threshold staining across the entorhinal cortex in **(A)** wild-type and *G2019S LRRK2* knockin mice, and **(B)** wild-type and *LRRK2* knockout mice. Data are expressed as bars depicting the mean ± SEM from either the ipsilateral or the contralateral hemisphere with each data point representing an animal (*n* = 7 animals/genotype). \*\**P*<0.01 by paired *t*-test within genotype, as indicated. Non-significant by unpaired *t*-test or Mann Whitney test between genotypes. **C)** Representative histological section displaying Iba1-positive microglia with accompanying image showing a generated HALO mask for low (yellow), intermediate (orange), and strong (red) area quantification thresholds. **D-E)** Quantification of the mean microglia cell body area in the entorhinal cortex in **(D)** wild-type and *G2019S LRRK2* knockin mice, and **(E)** wild-type and *LRRK2* knockout mice. Data are expressed as bars depicting the mean ± SEM from either the ipsilateral or the contralateral hemisphere with each data point representing an animal (*n* = 7 animals/genotype). \**P*<0.05, \*\**P*<0.01, or \*\*\**P*<0.001 by paired *t*-test within genotype, as indicated. \**P*<0.05 by unpaired *t*-test between genotypes. **F-G)** Histograms displaying the distribution of detected microglia cell body size in the ipsilateral and contralateral entorhinal cortices of **(F)** wild-type and *G2019S LRRK2* knockin mice, and **(G)** wild-type and *LRRK2* knockout mice. **H)** Representative histological sections depicting a “resting” microglia and an “activated” microglia with enlarged cell body, accompanied by generated HALO masks for microglial morphologic analysis.

Overall, this analysis provides clear indication that microglial activation in the entorhinal cortex, based on morphologic change, occurs in response to pathologic α-synuclein inclusions but is not drastically altered by LRRK2. We were not able to conduct a comparable analysis of morphologic change in the hippocampus due to substantially higher microglial density, limiting segmentation quality and identification of individual microglia. Qualitatively, we did observe morphologic changes in the hippocampus of PFF-injected *LRRK2* knockout mice, suggesting that the reduction in Iba1 immunostaining may be more related to microglial recruitment rather than activation, *per se*. It is also plausible that extracellular α-synuclein PFFs provide a more potent (or perhaps LRRK2-dependent) microglial stimuli relative to intraneuronal pathology, altering the inflammatory landscape within the hippocampus of *LRRK2* knockout mice.

### Pathological accumulation of **α**-synuclein inclusions across the hippocampal axis associates with neuropathologic changes independent of LRRK2

Neuropathologic changes involve a variety of concomitant pathologies along with reactive and compensatory modifications. Due to the static nature of histological assessment, the causal relationship between these features is not always clear. In our model, we have the advantage of initiating α-synuclein aggregation as a primary insult, through which we might understand how other changes are affected. We identified a clear gradient in α-synuclein pathology across the dorsal-ventral axis of the hippocampus in our model (**Figure 3A-D, 4A-D**). Thus, we sought to determine how this gradient in α-synuclein pathology was related to the burden of other markers of interest (**Figure 9A**). We found that these related neuropathological changes were tightly associated with the burden of α-synuclein pathology, suggesting that these secondary features were tightly linked to the burden of intraneuronal inclusions. We found that α-synuclein pathology strongly correlated with phosphorylated tau, GFAP, and Iba1 immunostaining in both wild-type mice as well as in G2019S LRRK2 knockin mice (**Figure 9B-D**). Likewise, significant correlations were also detected in both wild-type mice and *LRRK2* knockout mice in the other cohort (**Figure 9E-G**). Of note, α-synuclein pathology and Iba1 immunostaining did correlate significantly in *LRRK2* knockout mice. Thus, under the conditions of our model, we find that the level of intraneuronal pathology across the hippocampal axis was strongly associated with related markers, regardless of *LRRK2* genotype.

**Figure 9.**
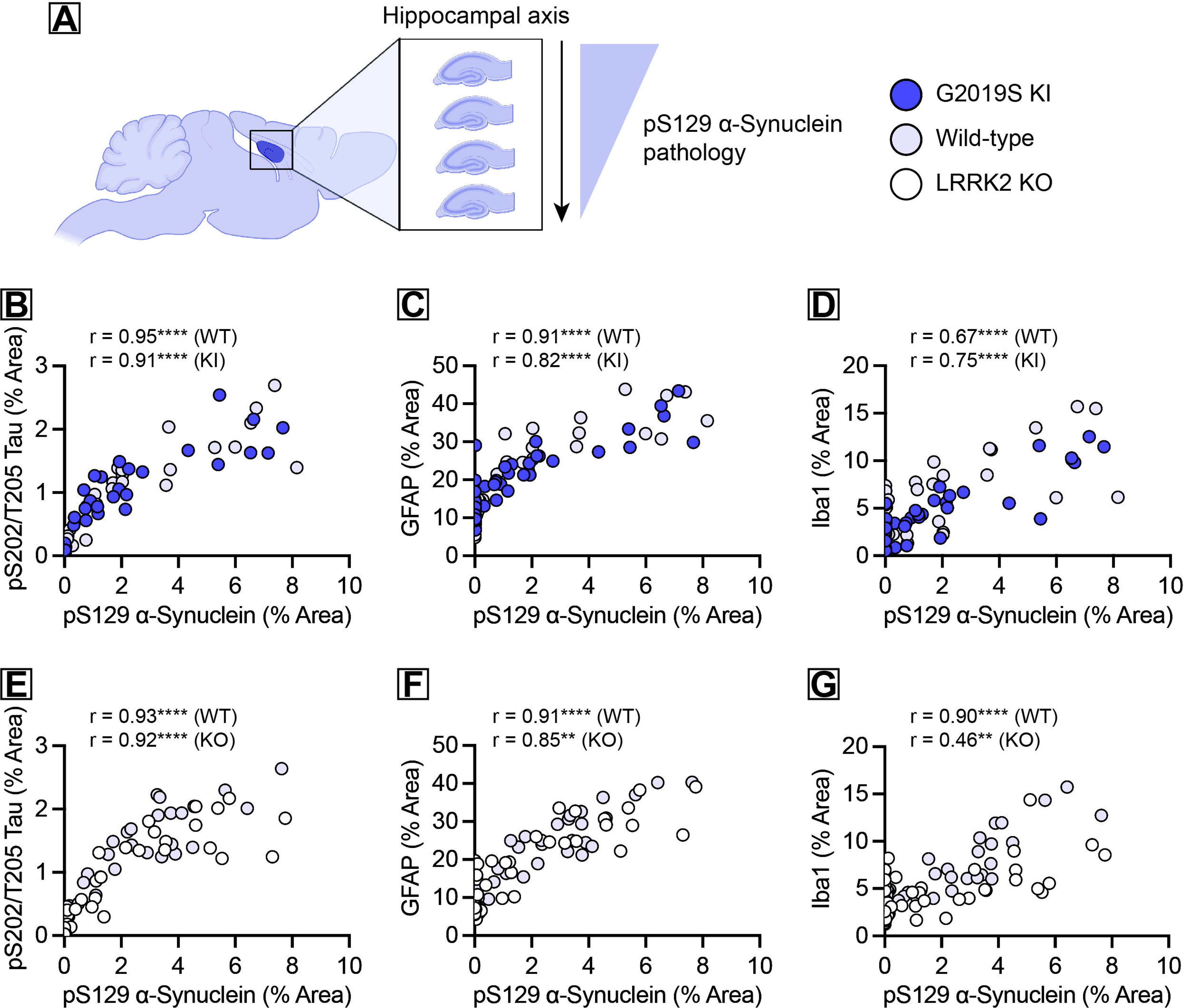
Pathological accumulation of α-synuclein inclusions across the hippocampal axis correlates with neuropathologic changes independent of *LRRK2* genotype. **A)** Schematic of the mouse hippocampal dorsal-ventral axis displaying serial sections with an observed α-synuclein pathology gradient. **B-D)** Association between % area quantification of pS129-α-synuclein immunostaining across the hippocampal axis and **(B)** pS202/T205-tau, **(C)** GFAP, and **(D)** Iba1 in wild-type and *G2019S LRRK2* knockin mice. \*\**\**\**P*<0.0001 by Spearman correlation, *r* values listed by genotype, as indicated. **E-G)** Association between % area quantification of pS129-α-synuclein immunostaining across the hippocampal axis and **(E)** pS202/T205-tau, **(F)** GFAP, and **(G)** Iba1 in wild-type and *LRRK2* knockout mice. \*\**P*<0.01 or \*\**\**\**P*<0.0001 by Spearman correlation, *r* values listed by genotype, as indicated.

## Discussion

Here, we characterize the influence of LRRK2 in a forebrain α-synuclein PFF model. We find that endogenous LRRK2 expression is not required for the templated formation of intraneuronal inclusions. Further, neither genetic deletion of *LRRK2* nor presence of the pathogenic G2019S *LRRK2* mutation had a substantial impact on the burden of forebrain α-synuclein pathology at 1 MPI. These findings build upon prior studies in transgenic α-synucleinopathy mouse models, where α-synuclein pathology was essentially independent of LRRK2 (Daher et al., 2012; Herzig et al., 2012). Consistent with our results, a mouse-based study utilizing α-synuclein viral vectors found no impact of *LRRK2* ablation on α-synuclein pathology but did detect a LRRK2-dependent effect on microglial activation (Perren et al., 2021). More relevant to the PFF model, pharmacologic inhibition of LRRK2 kinase activity was found to have little impact on the formation of pathology elicited by α-synuclein PFF both *in vitro* and *in vivo* (Henderson et al., 2018, 2019b).

Despite the lack of interaction in several paradigms, a number of studies have found a robust interaction between LRRK2 and α-synuclein. Antisense oligonucleotide (ASO)-mediated knockdown of *LRRK2* was demonstrated to dramatically reduce α-synuclein pathology *in vivo* (Zhao et al., 2017). Another study found that aged G2019S LRRK2 transgenic mice yielded elevated α-synuclein pathology in a viral vector model (Novello et al., 2018). Genetic modulation of LRRK2 in a transgenic α-synuclein mouse with extensive forebrain α-synuclein was shown to bidirectionally modulate α-synuclein pathology (Lin et al., 2009). Specific to the α-synuclein PFF model, G2019S LRRK2 BAC mice had a modest, but significant increase in α-synuclein inclusions under the intrastriatal paradigm (Bieri et al., 2019). It is unclear why various experimental models reveal such contrasting results, but several factors may be important to consider. Our study utilized the more recently developed G2019S LRRK2 knockin mice, which faithfully recapitulates endogenous LRRK2 expression, while previous studies utilized LRRK2 overexpression paradigms under various promoters. These previously detected interactions may be a feature of supraphysiologic LRRK2 expression, or the result of ectopic expression across various cell types. As well, different modalities may contain a significant inflammatory element, such as immunogenicity against viral vector components or PFF preparations. Given the relationship between LRRK2 and inflammation, it is possible that additional immune-activating components lead to profoundly different outcomes regarding α-synuclein pathology. Indeed, LRRK2-dependent inflammation may lead to incremental changes in α-synuclein pathology that become amplified over time.

Beyond inflammatory changes, LRRK2 is known to impact neurite development and alter neuronal biology which might affect the localization of endogenous α-synuclein (Brzozowski et al., 2021; MacLeod et al., 2006). It is possible that LRRK2 pathogenicity may lead to subtle changes that are masked in our model but may be detected in other experimental approaches. Towards this, a recent study administering α-synuclein PFFs to primary neurons derived from *LRRK2* knockout, wild-type, or G2019S LRRK2 knockin mice found a modest impact of both mutant LRRK2 and *LRRK2* knockout on the modulation of α-synuclein pathology (MacIsaac et al., 2020). It is possible that LRRK2 pathogenicity or loss of endogenous *LRRK2*, is less compensated for *in vitro*, leading to these results. Alternatively, greater variability in the *in vivo* model may mask more subtle effects that are detectable in primary neurons. Neuritic α-synuclein accumulation was especially influenced by *LRRK2* genotype in this study, so it is possible that our results are due to assessment at a stage beyond which LRRK2 might have an impact. Injection of a lower concentration of α-synuclein PFFs and assessment at an earlier timepoint might identify more immediate effects of LRRK2 on α-synuclein accumulation *in vivo*. We did not discriminate between neuritic and perisomatic α-synuclein pathology in our model, but it is important to consider that α-synuclein may be modulated in a compartment-specific manner.

Considering the approach of LRRK2 suppression may be informative. Examining the impact of conditional *LRRK2* knockdown in adult mice may provide some insight into these phenomena. Given the noteworthy reduction of α-synuclein pathology following ASO-mediated *LRRK2* knockdown, this approach is not without merit (Zhao et al., 2017). It is also plausible that the potential interaction of LRRK2 with α-synuclein is restricted to a particular neuronal population, such as dopaminergic neurons of the substantia nigra. We opted to examine the forebrain, as this region has high levels of endogenous LRRK2 expression and represents a key stage in the progression of Lewy-related pathology. Given the advent of LRRK2-targeted therapeutics for Parkinson’s disease, it is appropriate to determine whether a LRRK2-targeted approach might modify the progression of Lewy-related pathology in the forebrain with clinical relevance for Lewy-related cognitive dysfunction. Developmental compensation in *LRRK2* knockout mice may mask changes that are present in conditional models of *LRRK2* knockdown. Finally, given that age is the greatest risk factor for PD and *LRRK2*-PD typically presents with late-onset disease, the use of young adult mice may conceal age-dependent changes in LRRK2 biology.

Lewy-related pathology is a common incidental finding in the aged brain and in the absence of clinical manifestations (Adler et al., 2010). A pathogenic *LRRK2* mutation is not required for the formation of α-synuclein inclusions. In the case of *LRRK2*-PD, aberrant LRRK2 activity might modify the propagation of age-associated Lewy-related pathology, rather than contribute to its initial development or aspects of formation. While a variety of LRRK2 mouse models have been generated, none have convincingly revealed the *de novo* accumulation of α-synuclein. Given that many *LRRK2*-PD cases do not exhibit Lewy-related pathology, this concept of modification may be particularly relevant. While we did not detect a change in the initial formation of α-synuclein pathology, we limited our evaluation to 1 MPI. Given a longer period, it is plausible that *LRRK2* genotype might impact the pathologic propagation of α-synuclein via perturbation of the propagation process. In support of this hypothesis, a longitudinal study in G2019S LRRK2 BAC mice found subtle regional changes in the propagation of α-synuclein pathology over time (Henderson et al., 2019a). It is not clear if similar findings might be observed in G2019S LRRK2 knockin mice, or whether long-term LRRK2 inhibition might mitigate the spatiotemporal spread of α-synuclein pathology. A longitudinal study examining α-synuclein propagation over a longer duration is thus warranted in this model.

Our primary LRRK2-dependent finding within this forebrain model was the modest suppression of microglial activation in the dorsal hippocampus of *LRRK2* knockout mice relative to wild-type mice. Interestingly, we still observed activated microglial morphologies in both the hippocampus and entorhinal cortex of *LRRK2* knockout mice, suggesting that the impact of *LRRK2* deletion on microglial biology is likely complex and region or stimuli-specific. Indeed, previous work has demonstrated that LRRK2 directly alters microglial responses to α-synuclein PFFs (Moehle et al., 2012). The microglial response to intraneuronal inclusions is less amenable to *in vitro* study, but future efforts might provide clarity in the differential microglial response to fibrillar α-synuclein directly versus intraneuronal inclusions.

Given our findings, it appears unlikely that LRRK2 activity substantially influences the accumulation of α-synuclein pathology in a direct manner. With the extreme pathologic heterogeneity observed in *LRRK2*-PD and complete absence of Lewy-related pathology in some *LRRK2*-PD cases, it appears plausible that the detected neuropathologies in *LRRK2*-PD are the result of some common cellular dysfunction linked to LRRK2 pathobiology rather than an obligate manifestation. LRRK2 pathogenicity might facilitate a cascade in which α-synuclein aggregation is indirectly augmented, though it remains uncertain whether LRRK2 is a viable target for this aspect of disease. Collectively, these findings suggest that LRRK2-targeted therapies may need to be further evaluated in the context of Lewy-related pathology.

## Materials and Methods

### Animals

This study used G2019S LRRK2 knockin (homozygous KI/KI) and *LRRK2* knockout (homozygous KO/KO) mice and their wild-type littermates maintained on a C57BL/6J background. G2019S LRRK2 knockin mice carry two nucleotide substitutions in exon 41 of the *LRRK2* gene and were obtained from The Jackson Laboratory (B6.Cg-*Lrrk2^tm1.1Hlme^*/J, Strain # 030961). *LRRK2* knockout mice used in this study had a deletion of exon 41, as previously reported (Herzig et al., 2012). *LRRK2* knockout mice were kindly provided by Giorgio Rovelli and Derya Shimshek (Novartis Pharma AG, Basel Switzerland). All mice were housed under a 12-hour light/dark cycle and provided food *ad libitum*. Mice were maintained in accordance with NIH Guidelines for Care and Use of Laboratory Animals. Experiments were reviewed and approved by the Institutional Animal Care and Use Committee at Van Andel Institute.

### *In situ* hybridization and immunofluorescence

*In situ* RNA hybridization was performed using RNAScope Multiplex Fluorescent Detection kit v2 (Advanced Cell Diagnostics, Cat No. 323100) with the Akoya Biosciences Opal^TM^ fluorophores System (FP1488001KT) according to the manufacturer’s instructions. Briefly, fixed brains were flash frozen and subjected to 35-μm-thick horizontal sections using cryostat (Leica) and mounted on SuperFrost plus slides (Fisher Scientific). A probe recognizing mRNAs that encode mouse LRRK2 (Advanced Cell Diagnostics, Cat No. 421551) was used. Immunofluorescence were performed following *in situ* hybridization with anti-NeuN (MAB377, Millipore) and anti-Iba1 (019-19741, Wako) antibodies.

### Preparation and stereotactic injection of mouse **α**-synuclein PFFs

Recombinant mouse α-synuclein pre-formed fibrils (PFF) were generated and validated following previously established protocols (Dues et al., 2023; Volpicelli-Daley et al., 2014; Wang et al., 2019). Fibrils were diluted to a concentration of 5 μg/μl for *in vivo* use. Prior to injection, fibrils were thawed to room temperature and sonicated for 4 min using a Q700 cup horn sonicator (Qsonica) at 50% power for 120 pulses (1 s on, 1 s off). Mice were anesthetized with 2% isoflurane and positioned within a stereotactic frame (Kopf). Following incision, the skin flaps were retracted, and mice received a unilateral injection of mouse α-synuclein pre-formed fibrils using the following coordinates in relation to bregma: anterior-posterior (A-P), -2.5 mm; medio-lateral (M-L), +2.0 mm; dorso-ventral (D-V), -2.4, -1.4 mm. Each injection was delivered in a volume of 1 μl at a flow rate of 0.2 μl/min. All mice were sacrificed at 1-month post-injection.

### Immunohistochemistry

Whole brains were collected following transcardial perfusion with 4% paraformaldehyde (PFA) in 0.1 M phosphate buffer (PB) at physiologic pH. Brains were fixed for a 24-hour period in 4% PFA followed by cryoprotection in 30% sucrose in 0.1 M PB. Brains were cut into 35 μm-thick horizontal sections. Brain sections were quenched for endogenous peroxidase activity with 3% H_2_O_2_ (Sigma) in a methanol solution for 5 minutes at 4°C. Blocking was applied with 10% normal goat serum and 0.1% Triton-X100 in PBS for 1 hour at room temperature. Primary antibody incubation was completed for 48 hours at 4°C followed by incubation with biotinylated secondary antibodies (Vector Labs) for 24 hours at 4°C. Following incubation with ABC reagent (Vector Labs) for 1 hour at room temperature, sections were resolved in 3, 3’-diaminobenzidine tetrahydrochloride (DAB; Vector Labs). Immunostained sections were subsequently mounted on Superfrost plus slides (Fisher Scientific), dehydrated with increasing ethanol concentrations and xylene, and coverslipped using Entellan mounting medium (Merck). The following primary antibodies were used: rabbit anti-pS129-α-synuclein (ab51253; Abcam), mouse anti-pS202/T205-tau (MN1020, Thermo Fisher), rabbit anti-Iba1 (019-19741, Waco), mouse anti-GFAP (G3893, Sigma). The following biotinylated secondary antibodies were used: goat anti-mouse IgG and goat anti-rabbit IgG (Vector Labs).

### Digital and quantitative pathology

Slide-mounted horizontal brain sections were scanned using a ScanScope XT slide scanner (Aperio) at a resolution of 0.5 µm/pixel. Annotation and quantitative assessment of the hippocampal and cortical regions of interest (6 complete horizontal brain sections assessed per animal, left/right hemispheric regions were annotated and assessed separately) were performed using HALO analysis software (Area quantification, Object colocalization, and Microglial activation modules; Indica Labs Inc.). For each quantitative approach, threshold parameters were developed and optimized to detect positive-staining without background staining or tissue-level artifacts. All sections/slides were assessed using the optimized HALO batch analysis function for each marker of interest. All annotations and assessments were conducted agnostic to slide identification or animal genotype.

### Statistical Analyses

Statistical analysis of all data was performed with GraphPad Prism 9 (GraphPad Software). Bar graphs depict all data points with the mean ± SEM. The data was checked for Gaussian distribution using the Shapiro-Wilk normality test. Statistical significance within genotype (ipsilateral vs contralateral) was assessed by either paired *t*-test or Wilcoxon matched-pairs signed rank test. Statistical significance between genotype (WT vs G2019S KI or WT vs LRRK2 KO) was assessed by either unpaired *t*-test or Mann Whitney test. Graphs were generated within GraphPad Prism.

## Funding

The authors are appreciative for financial support for this work from the National Institutes of Health (R56 AG074473 to D.J.M.) and the Van Andel Institute Graduate School.

## Acknowledgements

We acknowledge the VAI Vivarium, and Pathology and Biorepository Cores for technical assistance. Some figures were developed using graphic design tools in BioRender (Biorender.com).

## Notes

### Competing Interest Statement

The authors have declared no competing interest.

