## Supplemental Data for "Formation of templated inclusions in a forebrain α-synuclein mouse model is independent of LRRK2"

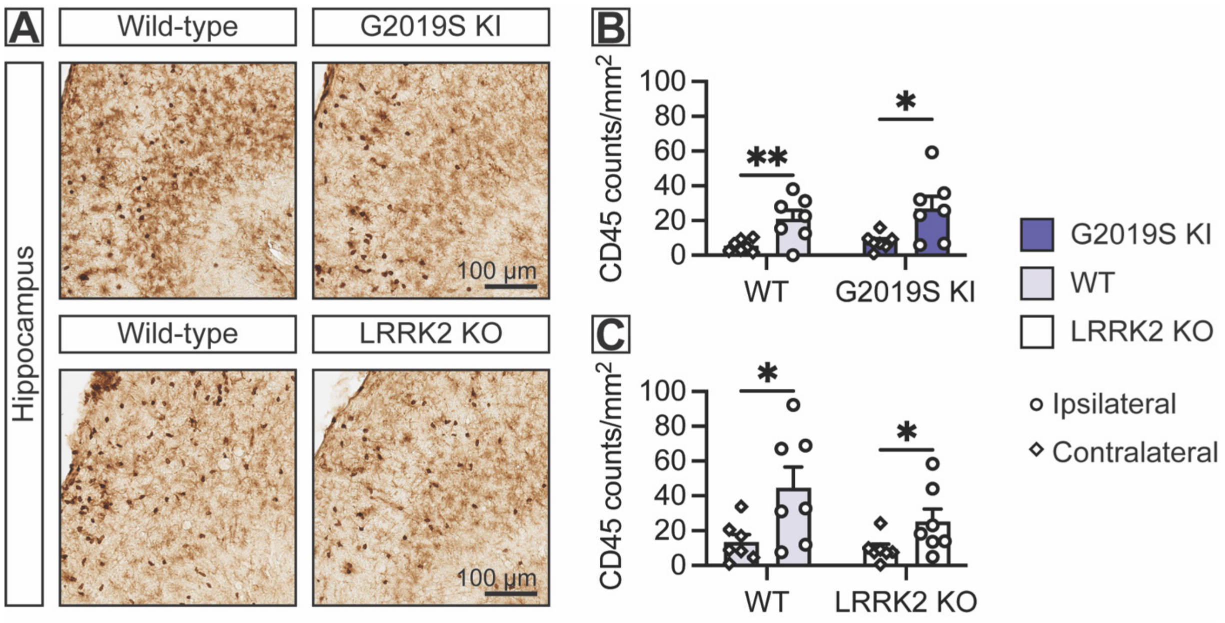


**Supplemental Figure 1. CD45-positive peripheral immune cell recruitment to the hippocampus following injection with α-synuclein PFFs is not significantly altered by *LRRK2*.**

**A)** Representative histological sections of the dorsal hippocampus immunostained for CD45 in wild-type controls and *G2019S LRRK2* knockin and *LRRK2* knockout mice. Scale bars: 100 µm. **B)** Quantification of CD45-positive cell counts/mm$^2$ in wild-type control and *G2019S LRRK2* knockin mice ($n$ = 7 animals/genotype). **C)** Quantification of CD45-positive cell counts/mm$^2$ in wild-type control and *LRRK2* knockout mice ($n$ = 7 animals/genotype). Data are expressed as bars depicting the mean ± SEM from either the ipsilateral (injected) hemisphere or the contralateral (non-injected) hemisphere with each data point representing an animal. *$P$<0.05 or **$P$<0.01 by paired $t$-test or Wilcoxon matched-pairs signed rank test within genotype, as indicated. Non-significant by unpaired $t$-test or Mann Whitney test between genotypes.
